# Gene model for the ortholog of *tgo* in *Drosophila pseudoobscura*

**DOI:** 10.1101/2025.07.13.664599

**Authors:** Megan E. Lawson, Janelle Marie Suriaga, Mary B. Rowland, Laura K. Reed, Chinmay P. Rele, Jacqueline K. Wittke-Thompson

## Abstract

Gene model for the ortholog of *tango* (*tgo*) in the April 2013 (BCM-HGSC Dpse_3.0/DpseGB3) Genome Assembly (GenBank Accession: GCA_000001765.2) of *Drosophila pseudoobscura*. This ortholog was characterized as part of a developing dataset to study the evolution of the Insulin/insulin-like growth factor signaling pathway (IIS) across the genus *Drosophila* using the Genomics Education Partnership gene annotation protocol for Course-based Undergraduate Research Experiences.

## Introduction

*This article reports a predicted gene model generated by undergraduate work using a structured gene model annotation protocol defined by the Genomics Education Partnership (GEP; thegep.org) for Course-based Undergraduate Research Experience (CURE). The following information in quotes may be repeated in other articles submitted by participants using the same GEP CURE protocol for annotating Drosophila species orthologs of Drosophila melanogaster genes in the insulin signaling pathway*.

“In this GEP CURE protocol students use web-based tools to manually annotate genes in non-model *Drosophila* species based on orthology to genes in the well-annotated model organism fruit fly *Drosophila melanogaster*. The GEP uses web-based tools to allow undergraduates to participate in course-based research by generating manual annotations of genes in non-model species (Rele et al., 2023). Computational-based gene predictions in any organism are often improved by careful manual annotation and curation, allowing for more accurate analyses of gene and genome evolution (Mudge and Harrow 2016; Tello-Ruiz et al., 2019). These models of orthologous genes across species, such as the one presented here, then provide a reliable basis for further evolutionary genomic analyses when made available to the scientific community.” (Myers et al., 2024).

“The particular gene ortholog described here was characterized as part of a developing dataset to study the evolution of the Insulin/insulin-like growth factor signaling pathway (IIS) across the genus *Drosophila*. The Insulin/insulin-like growth factor signaling pathway (IIS) is a highly conserved signaling pathway in animals and is central to mediating organismal responses to nutrients (Hietakangas and Cohen 2009; Grewal 2009).” (Myers et al., 2024).

“The Drosophila *tango* (CG11987, FBgn0264075) gene encodes a bHLH-PAS protein that controls CNS midline and tracheal development (Sonnenfeld et al., 1997). Both cell culture and in vivo studies have shown that a DNA enhancer element acts as a binding site for both Single-minded::Tango and Trachealess::Tango heterodimers and functions in controlling CNS midline and tracheal transcription. Isolation and analysis of tango mutants reveal CNS midline and tracheal defects. In addition, the bHLH-PAS proteins Similar (Sima) and Tango (Tgo) function as HIF-α and HIF-β homologues, respectively, in a conserved hypoxia-inducible transcriptional response in *Drosophila melanogaster* that is homologous to the mammalian HIF-dependent response (Lavista-Llanos et al., 2002). Insulin has been shown to activate HIF-dependent transcription, both in *Drosophila* S2 cells and in living *Drosophila* embryos, and this effect is mediated by PI3K-AKT and TOR pathways. Overexpression of dAKT and dPDK1 in normoxic embryos provoked a major increase in Sima nuclear localization, mimicking the effect of a hypoxic treatment (Dekanty et al., 2005).” (Lawson 2025a).

“*D. pseudoobscura* is part of the *pseudoobscura* species subgroup within the *obscura* species group in the subgenus *Sophophora* of the genus *Drosophila* (Sturtevant 1942; Buzzati-Traverso & Scossiroli 1955). It was first described by Frolowa (1929). The *pseudoobscura* species subgroup is endemic to the New World, where *D. pseudoobscura* is distributed throughout Western North American, and south through Mexico and Central America (Markow & O’Grady 2005). An additional population of *D. pseudoobscura*, found near Bogota, Columbia, is partially reproductively isolated from the North and Central American populations (Prakash 1972). It is found primarily in chaparral and temperate forests. *D. pseudoobscura* has been studied extensively in the context of ecological and behavioral genetics, speciation, and genome evolution (Powell 1997; Lawson et al., 2024).” (Lawson 2025b).

We propose a gene model for the ortholog in *D. pseudoobscura* (NCBI taxon ID 46245) of the *D. melanogaster* tango (*tgo*) gene. The genomic region of the ortholog corresponds to the uncharacterized protein XP_001359833.1 (Locus ID LOC4803031) in the Apr. 2013 (BCM-HGSC Dpse_3.0/DpseGB3) Genome Assembly of *D. pseudoobscura* (GCA_000001765.2, Chen et al., 2014). This model is based on RNA-Seq data from *D. pseudoobscura* (SRP006203) and *tgo* in *D. melanogaster* using FlyBase release FB2024_02 (GCA_000001215.4; Gramates et al., 2022; Jenkins et al., 2022; Larkin et al., 2021).

### Synteny

The target gene, *tgo*, occurs on chromosome 3R in *D. melanogaster* and is flanked upstream by *neuralized* (*neur*) and *hyrax* (*hyx*) and downstream by *CG11986* and *Splicing factor 3b subunit 5* (*Sf3b5*). The *tblastn* search of *D. melanogaster* tgo-PA (query) against the *D. pseudoobscura* (GenBank Accession: GCA_000001765.2 Genome Assembly (database) placed the putative ortholog of *tgo* within scaffold CM000070 (CM000070.4) at locus LOC4803031 (XP_001359833.1)— with an E-value of 0.0 and a percent identity of 83.05%. Furthermore, the putative ortholog is flanked upstream by LOC4803030 (XP_001359832.3) and LOC4803029 (XP_001359831.1), which correspond to *neur* and *hyx* in *D. melanogaster* (E-value: 0.0 and 0.0; identity: 86.62% and 91.48%, respectively, as determined by *blastp*; Figure 1A; Altschul et al., 1990). The putative ortholog of *tgo* is flanked downstream by LOC4803033 (XP_033232624.1) and LOC4803032 (XP_015038250.1), which correspond to *CG11986* and *Kcmf1* in *D. melanogaster* (E-value: 0.0 and 0.0; identity: 81.39% and 81.19%, respectively, as determined by *blastp*). The putative ortholog assignment for *tgo* in *D. pseudoobscura* is supported by the following evidence: Most of the genes surrounding the *tgo* ortholog are orthologous to the genes at the same locus in *D. melanogaster* and local synteny is mostly conserved aside from the second most downstream gene, supported by E-values and percent identities, so we conclude that LOC4803031 is the correct ortholog of *tgo* in *D. pseudoobscura* (Figure 1A).

**Figure 1.**
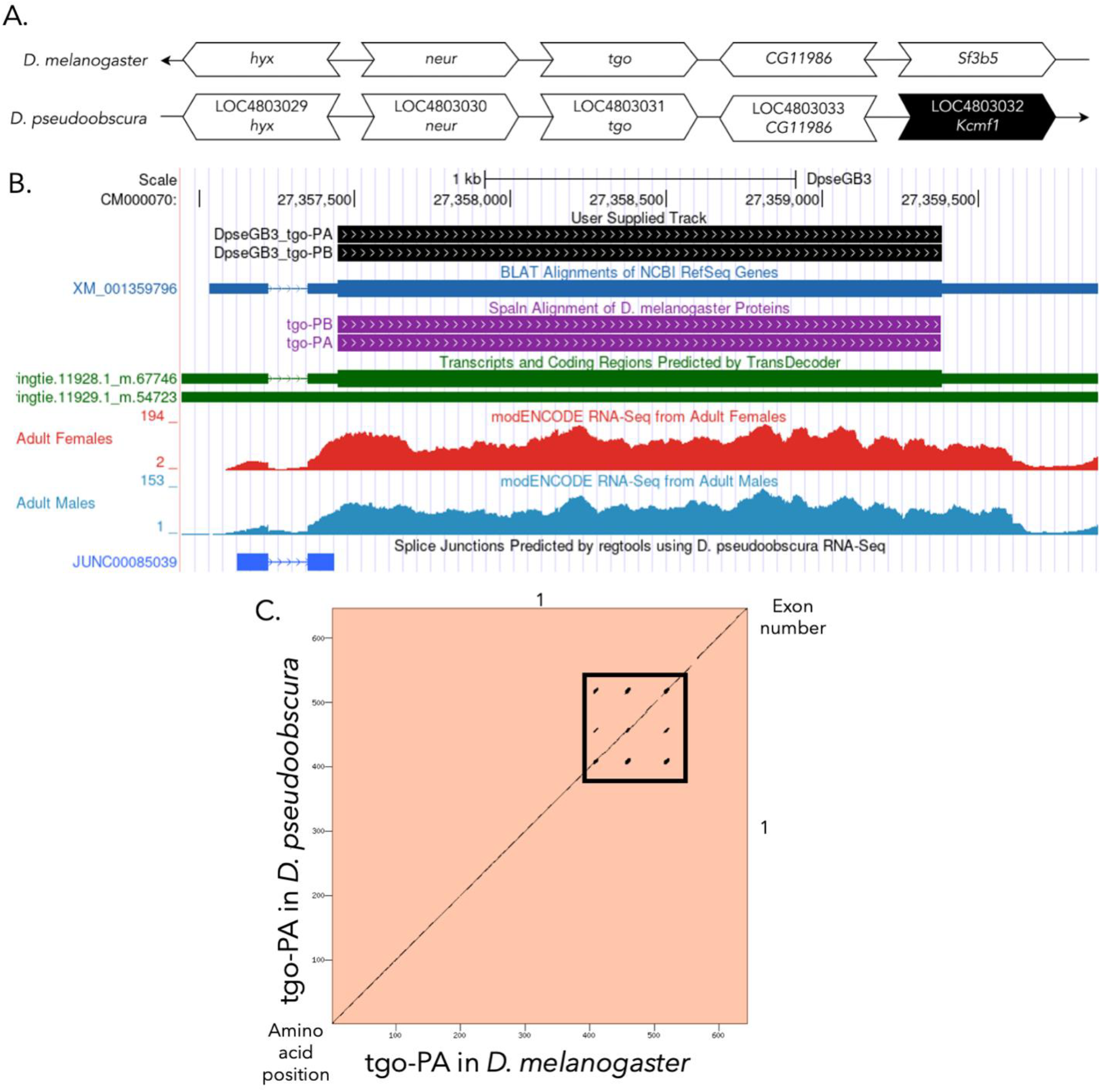
*tgo* gene model comparison between *Drosophila pseudoobscura* and *Drosophila melanogaster* orthologs. **(A) Synteny comparison of the genomic neighborhoods for *tgo* in *Drosophila melanogaster* and *D. pseudoobscura***. Thin underlying arrows indicate the DNA strand within which the target gene–*tgo*–is located in *D. melanogaster* (top) and *D. pseudoobscura* (bottom). The thin arrow pointing to the left indicates that *tgo* is on the negative (-) strand in *D. melanogaster*, and the thin arrow pointing to the right indicates that *tgo* is on the positive (+) strand in *D. pseudoobscura*. The wide gene arrows pointing in the same direction as *tgo* are on the same strand relative to the thin underlying arrows, while wide gene arrows pointing in the opposite direction of *tgo* are on the opposite strand relative to the thin underlying arrows. White gene arrows in *D. pseudoobscura* indicate orthology to the corresponding gene in *D. melanogaster*, while black gene arrows indicate non-orthology. Gene symbols given in the *D. pseudoobscura* gene arrows indicate the orthologous gene in *D. melanogaster*, while the locus identifiers are specific to *D. pseudoobscura*. **(B) Gene Model in GEP UCSC Track Data Hub** (Raney et al., 2014). The coding-regions of *tgo* in *D. pseudoobscura* are displayed in the User Supplied Track (black); coding exons are depicted by thick rectangles and introns by thin lines with arrows indicating the direction of transcription. Subsequent evidence tracks include BLAT Alignments of NCBI RefSeq Genes (dark blue, alignment of Ref-Seq genes for *D. pseudoobscura*), Spaln of D. melanogaster Proteins (purple, alignment of Ref-Seq proteins from *D. melanogaster*), Transcripts and Coding Regions Predicted by TransDecoder (dark green), RNA-Seq from Adult Females and Adult Males (red and light blue, respectively; alignment of Illumina RNA-Seq reads from *D. pseudoobscura*), and Splice Junctions Predicted by regtools using *D. pseudoobscura* RNA-Seq (SRP006203). Splice junctions shown have a minimum read-depth of 10 with 10-49 supporting reads indicated in blue. **(C) Dot Plot of tgo-PA in *D. melanogaster* (*x*-axis) vs. the orthologous peptide in *D. pseudoobscura* (*y*-axis)**. Amino acid number is indicated along the left and bottom; coding-exon number is indicated along the top and right, and exons are also highlighted with alternating colors. Line breaks in the dot plot indicate mismatching amino acids at the specified location between species. The boxed region highlights a tandem repeat within the amino acid sequence.

### Protein Model

*tgo* in *D. pseudoobscura* has two identical protein-coding isoforms (tgo-PA and tgo-PB; Figure 1B). These isoforms (tgo-PA and tgo-PB) contain one protein-coding exon. Relative to the ortholog in *D. melanogaster*, the coding-exon number and isoform count are conserved, as *tgo* in *D. melanogaster* also has two identical isoforms, each with one exon. The sequence of tgo-PA in *D. pseudoobscura* has 94.92% identity (E-value: 0.0 with the protein-coding isoform tgo-PA in *D. melanogaster*, as determined by *blastp* (Figure 1C). Coordinates of this curated gene model are stored by NCBI at GenBank/BankIt (accessions **<u>BK065289</u>** and **<u>BK065290</u>**). This gene model can also be seen within the target genome at this [TrackHub](https://gander.wustl.edu/cgi-bin/hgTracks?db=DpseGB3&lastVirtModeType=default&lastVirtModeExtraState=&virtModeType=default&virtMode=0&nonVirtPosition=&position=CM000070%3A27356945%2D27359885&hgsid=16589931_AE4EsfVcZH1UPckeql5I12UE9gMt).

### Methods

“Detailed methods including algorithms, database versions, and citations for the complete annotation process can be found in Rele et al. (2023). Briefly, students use the GEP instance of the UCSC Genome Browser v.435 (https://gander.wustl.edu; Kent WJ et al., 2002; Navarro Gonzalez et al., 2021) to examine the genomic neighborhood of their reference IIS gene in the *D. melanogaster* genome assembly (Aug. 2014; BDGP Release 6 + ISO1 MT/dm6). Students then retrieve the protein sequence for the *D. melanogaster* reference gene for a given isoform and run it using *tblastn* against their target *Drosophila* species genome assembly on the NCBI BLAST server (https://blast.ncbi.nlm.nih.gov/Blast.cgi; Altschul et al., 1990) to identify potential orthologs. To validate the potential ortholog, students compare the local genomic neighborhood of their potential ortholog with the genomic neighborhood of their reference gene in *D. melanogaster*. This local synteny analysis includes at minimum the two upstream and downstream genes relative to their putative ortholog. They also explore other sets of genomic evidence using multiple alignment tracks in the Genome Browser, including BLAT alignments of RefSeq Genes, Spaln alignment of *D. melanogaster* proteins, multiple gene prediction tracks (e.g., GeMoMa, Geneid, Augustus), and modENCODE RNA-Seq from the target species. Detailed explanation of how these lines of genomic evidenced are leveraged by students in gene model development are described in Rele et al. (2023). Genomic structure information (e.g., CDSs, intron-exon number and boundaries, number of isoforms) for the *D. melanogaster* reference gene is retrieved through the Gene Record Finder (https://gander.wustl.edu/~wilson/dmelgenerecord/index.html; Rele et al., 2023).

Approximate splice sites within the target gene are determined using *tblastn* using the CDSs from the *D. melanogaste*r reference gene. Coordinates of CDSs are then refined by examining aligned modENCODE RNA-Seq data, and by applying paradigms of molecular biology such as identifying canonical splice site sequences and ensuring the maintenance of an open reading frame across hypothesized splice sites. Students then confirm the biological validity of their target gene model using the Gene Model Checker (https://gander.wustl.edu/~wilson/dmelgenerecord/index.html; Rele et al., 2023), which compares the structure and translated sequence from their hypothesized target gene model against the *D. melanogaster* reference gene model. At least two independent models for a gene are generated by students under mentorship of their faculty course instructors. Those models are then reconciled by a third independent researcher mentored by the project leaders to produce the final model. Note: comparison of 5’ and 3’ UTR sequence information is not included in this GEP CURE protocol.” (Gruys et al., 2025)

## Supporting information

Gene model data files

## Supplemental Files

1. Zip file containing a FASTA, PEP, GFF files for the gene model
2. Figure 1 in high resolution

**Metadata:** Bioinformatics, Genomics, *Drosophila*, Genotype Data, New Finding

## Acknowledgements

We would like to thank Wilson Leung for developing and maintaining the technological infrastructure that was used to create this gene model and Laura K. Reed for overseeing the project. Thank you to [FlyBase](https://flybase.org/) for providing the definitive database for *Drosophila melanogaster* gene models. Further, we would like to thank the editors and developers at the journal *microPublication: Biology* for assistance in developing the template for these single gene ortholog publications.

## References

Altschul SF, Gish W, Miller W, Myers EW, Lipman DJ. 1990. Basic local alignment search tool. J Mol Biol 215(3):403–410. PMID: 2231712.

Buzzati-Traverso AA, Scossiroli RE. 1955. The obscura group of the genus Drosophila. Adv Genet 7:47–92. PMID: 13258372

Chen ZX, Sturgill D, Qu J, Jiang H, Park S, Boley N, Suzuki AM, Fletcher AR, Plachetzki DC, FitzGerald PC, et al. 2014. Comparative validation of the D. melanogaster modENCODE transcriptome annotation. Genome Res 24(7):1209–1223. PMID: 24985915

Dekanty A, Lavista-Llanos S, Irisarri M, Oldham S, Wappner P. 2005. The insulin-PI3K/TOR pathway induces a HIF-dependent transcriptional response in Drosophila by promoting nuclear localization of HIF-α/Sima. J Cell Sci 118(23): 5431–5441. PMID: 16278294

Frolowa SL, Astaurow BL. 1929. Die Chromosomengarnitur als systematisches Merkmal. Z.Zellforsch 10: 201–213. 10.1007/BF02450642

Gramates LS, Agapite J, Attrill H, Calvi BR, Crosby M, dos Santos G Goodman JL, Goutte-Gattat D, Jenkins V, Kaufman T, Larkin A, Matthews B, Millburn G, Strelets VB, and the FlyBase Consortium. 2022. FlyBase: a guided tour of highlighted features. Genetics 220(4):iyac035. PMID: 35266522

Grewal SS. Insulin/TOR signaling in growth and homeostasis: a view from the fly world. 2009. Int J Biochem Cell Biol 41(5):1006–1010. PMID: 18992839

Gruys ML, Sharp MA, Lill Z, Xiong C, Hark AT, Youngblom JJ, Rele CP, Reed LK. 2025. Gene model for the ortholog of Glys in Drosophila simulans. microPubl Biol:10.17912/micropub.biology.001168. PMID: 39845267

Hietakangas V, Cohen SM. Regulation of tissue growth through nutrient sensing. 2009. Annu Rev Genet 43:389–410. PMID: 19694515

Jenkins VK, Larkin A, Thurmond J, FlyBase Consortium. 2022. Using FlyBase: A Database of Drosophila Genes and Genetics. Methods Mol Biol 2540:1–34. PMID: 35980571

Kent WJ, Sugnet CW, Furey TS, Roskin KM, Pringle TH, Zahler AM, Haussler D. 2002. The Human Genome Browser at UCSC. Genome Res 12:996–1006. PMID: 12045153

Larkin A, Marygold SJ, Antonazzo G, Attrill H, dos Santos G, Garapati PV, Goodman JL, Gramates LS, Millburn G, Strelets VB, Tabone CJ, and Thurmond J and the FlyBase Consortium. 2021. FlyBase: updates to the Drosophila melanogaster knowledge base. Nucleic Acids Res 49(D1): D899–D907. PMID: 33219682

Lavista-Llanos S, Centanin L, Irisarri M, Russo DM, Gleadle JM, Bocca SN, Mussopappa M, Ratcliffe PJ, Wappner P. 2002. Control of the hypoxic response in Drosophila melanogaster by the basic helix-loop-helix PAS protein similar. Mol Cell Biol 22(19):6842–6853. PMID: 12215541

Lawson ME, Dela Cruz M, Harrington DL, Vincent JA, McKenna C, Goodman A, Barnard D, Rele CP. 2024. Gene model for the ortholog of Pten in Drosophila miranda. microPubl Biol:10.17912/micropub.biology.000986. PMID: 39381638

Lawson M.E., Anderton Jr. RM, Perez J, Reed LK, Wittke-Thompson J, Rele CP. 2025. Gene model for the ortholog of tgo in Drosophila mojavensis. Manuscript submitted.

Lawson M.E., Dufur R, Wright B, Wellik IG, Long LJ, Thompson JS, Rele CP. 2025. Gene model for the ortholog of Roc1a in Drosophila pseudoobscura. Manuscript submitted.

Markow TA, O’Grady P. 2005. Drosophila: A guide to species identification and use. Academic Press, Amsterdam. 978-0-12-473052-6

Mudge JM, Harrow J. 2016. The state of play in higher eukaryote gene annotation. Nat Rev Genet 17:758–772. PMID: 27773922

Myers A, Hoffman A, Natysin M, Arsham AM, Stamm J, Thompson JS, Rele CP, Reed LK. 2024. Gene model for the ortholog Myc in Drosophila ananassae. microPubl Biol:10.17912/micropub.biology.000856. PMID: 39677519

Navarro Gonzalez J, Zweig AS, Speir ML, Schmelter D, Rosenbloom KR, Raney BJ, Powell CC, Nassar LR, Maulding ND, Lee CM et al. 2021. The UCSC Genome Browser database: 2021 update. Nucleic Acids Res 49(1):1046–1057. PMID: 33221922

Powell, JR. 1997. Progress and prospects in evolutionary biology: the Drosophila model. New York, NY: Oxford Academic Press. ISBN: 9780195076912

Prakash S. 1972. Origin of reproductive isolation in the absence of apparent genic differentiation in a geographic isolate of Drosophila pseudoobscura. Genetics 72(1):143–155. PMID: 5073854

Raney BJ, Dreszer TR, Barber GP, Clawson H, Fujita PA, Wang T, Nguyen N, Paten B, Zweig AS, Karolchik D, Kent WJ. 2014. Track data hubs enable visualization of user-defined genome-wide annotations on the UCSC Genome Browser. Bioinformatics 30(7):1003–1005. PMID: 24227676

Rele CP, Sandlin KM, Leung W, Reed LK. 2023. Manual annotation of Drosophila genes: a Genomics Education Partnership protocol. F1000Research 11:1579. https://doi.org/10.12688/f1000research.126839.2

Sonnenfeld M, Ward M, Nystrom G, Mosher J, Stahl S, Crews S. 1997. The Drosophila tango gene encodes a bHLH-PAS protein that is orthologos to mammalian Arnt and controls CNS midline and tracheal development. Development 124(22):4571–4582.

Sturtevant AH. 1942. The classification of the genus Drosophila with the description of nine new species. Univ Texas Publ 4213:5–51.

Tello-Ruiz MK, Marco CF, Hsu FM, Khangura RS, Qiao P, Sapkota S, Stitzer MC, Wasikowski R, Wu H, Zhan J et al. 2019. Double triage to identify poorly annotated genes in maize: The missing link in community curation. PLoS One 14:e0224086– e0224013. PMID: 31658277

